# Submicron Crystals of the Parkinson’s Disease *Substantia nigra*: Calcium Oxalate, Titanium Dioxide and Iron Oxide

**DOI:** 10.1101/523878

**Authors:** Adam Heller, Sheryl S. Coffman

## Abstract

Parkinson’s disease (PD) results of the death of dopaminergic neurons of the *substantia nigra.* When activated, the NLRP3 inflammasome of phagocytes releases inflammatory agents, their release resulting in the death of proximal cells. The hallmark protein of PD, aggregated α-synuclein, is phagocytized and activates the NLRP3 inflammasome. Because crystalline particles are known to activate the NLRP3 inflammasome, to enhance α-synuclein expression and aggregation in dopaminergic neurons and because their facets may mis-template adsorbed α-synuclein, we probe here, by transmission electron microscopy (TEM), four human PD *substantia nigra* specimens for their crystalline particles. Samples weighing 5 mg of PD stages 1, 2, 4 and 5 were processed by proteolysis and centrifugation. TEM-grids were dipped in the centrifugate diluted to 1 mL and the dried films were searched for crystalline particles. Two types of crystalline particles, known to activate the NLRP3 inflammasome were found. Endogenous calcium oxalate, a downstream product of ascorbate and dopamine oxidation-produced hydrogen peroxide; and TiO_2_, the with pigment of foods and medications. The number-density of the NLRP-inflammasome activating crystalline particles found approached the reported about-equal number-densities of microglia and neuronal cells in the brain.

The observations of COD and protein-coated TiO_2_ support two putative feedback loops, both leading to dopaminergic neuron death. In one, polymeric oxidized-dopamine catalyst accelerates H_2_O_2_-generation, the H_2_O_2_ indirectly oxidizing ascorbate in an ascorbate-fueled, oxalate-generating, loop the excess oxalate precipitating the subsequently inflammasome-activating COD crystals; In the second, protein-adsorbing facets of TiO_2_ mis-template the aggregation of α-synuclein to produce inflammasome-activating mis-folded α-synuclein.

## Introduction

Parkinson’s disease (PD) results of the degeneration of dopaminergic neurons of the *substantia nigra.* Studies of the past 5 years have shown that a cause of the degeneration is inflammatory attack of dopaminergic neurons of the *substantia nigra* by microglia when their NLRP3 inflammasome is activated by phagocytized misfolded α-synuclein. [1–12] The misfolded β-sheet-rich α-synuclein aggregates in PD’s hallmark Lewy bodies. If misfolded α-synuclein templates the misfolding of more synuclein, it constitutes a PD-propagating prion.[13–15] Because phagocytized crystals have been shown to activate the NLRP3 inflammasome [16–25], because they enhance in dopaminergic neurons α-synuclein expression and aggregation, [26] and they adsorb and orient peptides and as well as proteins,[27–29] we probe here by TEM if there are in the human PD *substantia nigra* crystals. We find ghosts of hydrated calcium oxalate (COD) and protein-coated titanium dioxide (TiO_2_) crystals. COD is the only non-phosphate containing crystalline insoluble calcium compound in mammalian tissues. It is a long-known cytotoxic component of urinary tract and kidney calculi, [30–32] associated also with retinopathy.[33, 34] Both COD and TiO_2_ are phagocytized, [25, 35–39] TiO_2_ also by microglia. [40] Both activate the NLRP3 inflammasome. [24, 25, 38, 41–47] Crystalline COD is associated with human lesions [31, 33, 34, 48–50] and has been reported in the human brain, [51] particularly when patients are lethally poisoned by the antifreeze ethylene glycol.[52–54] Ethylene glycol is metabolized to oxalic acid [55, 56] and is precipitated in the brain as calcium oxalate [57]. Ethylene glycol poisoning has induced PD symptoms in a patient.[58]

TiO_2_ is the dominant white pigment of foods and medications. It has been detected in brains of animals ingesting or inhaling the submicron-diameter pigment particles.[35, 59–61] In zebrafish larvae submicron TiO_2_ particles increased the expressions of PD’s hallmark Lewy body formation-associated*pink1, parkin, α-syn* and *uchl1* and caused loss of dopaminergic neurons. [62] Treatment of dopaminergic neurons with TiO_2_ increased their α-synuclein expression and caused aggregation of α-synuclein in a dose-dependent manner. [26] TiO_2_ adsorbs and orients peptides and proteins. [27–29]

### Materials and Methods

Four *substantia nigra* containing specimens of PD stages 1, 2, 4 and 5 were received from the NIH Harvard NeuroBioBank. Of each, 5 mg of each was excised and processed for TEM in plastic-ware free of crystalline matter. The minced samples were digested overnight at ambient temperature with 0.5 mL of an aqueous solution of 5 mg of trypsin (from bovine pancreas, Sigma catalog number 1426) and 5 mg of proteinase K (from *Tritirachium album*, Sigma catalog number P2308). The digested suspensions were centrifuged at 12,000g for 1 hour then the precipitate, with 0.5 mL added water, was dispersed in the resulting 1 mL volume. A 3-mm diameter, 314 square, 200 mesh, copper TEM grid with type-B carbon support (Ted Pella Redding CA, catalog number 1810) was dipped in the resulting 1 mL homogeneous dispersion and dried. The dipping and drying resulted in a residual protein, DNA and other biological matter containing film of a thickness that was about similar to that when a ~10 μL droplet was applied to the 3 mm 314-square grid. To rule out lab-ware-, water-and reagent-associated crystal contamination, samples without tissue were identically processed and were applied to TEM grids. No crystalline particles were observed.

A JEOL 2010F Transmission Electron Microscope (TEM) was used for imaging the particles. Their crystallinity was confirmed by their electron diffraction or by their energy-dispersive X-ray spectroscopic elemental analysis showing > 40 weight % of their metallic element. The TEM search protocol involved location darker, higher atomic-mass, and sharp-edged objects and elemental assay of their selected areas by energy dispersive X-ray spectroscopy and probing for electron diffraction by their crystalline lattices. The typical TEM time required to locate and characterize one crystalline particle was 2-3 hours. Squares of the 314-square grid were searched until 6 or more crystalline particles were found. In the grids of the PD stages 1 and 2 specimens, search of 3 squares was required; one square of the PD stage 4 grid was searched; and because the first PD stage 5 square searched had numerous particles (Figure 1) only part of it was searched. The apparent increase un the number of crystalline particles per square with the PD stage is likely misleading, because the distribution of crystalline particles on the dipped and dried TEM grid can be non-uniform.

**Figure 1.**
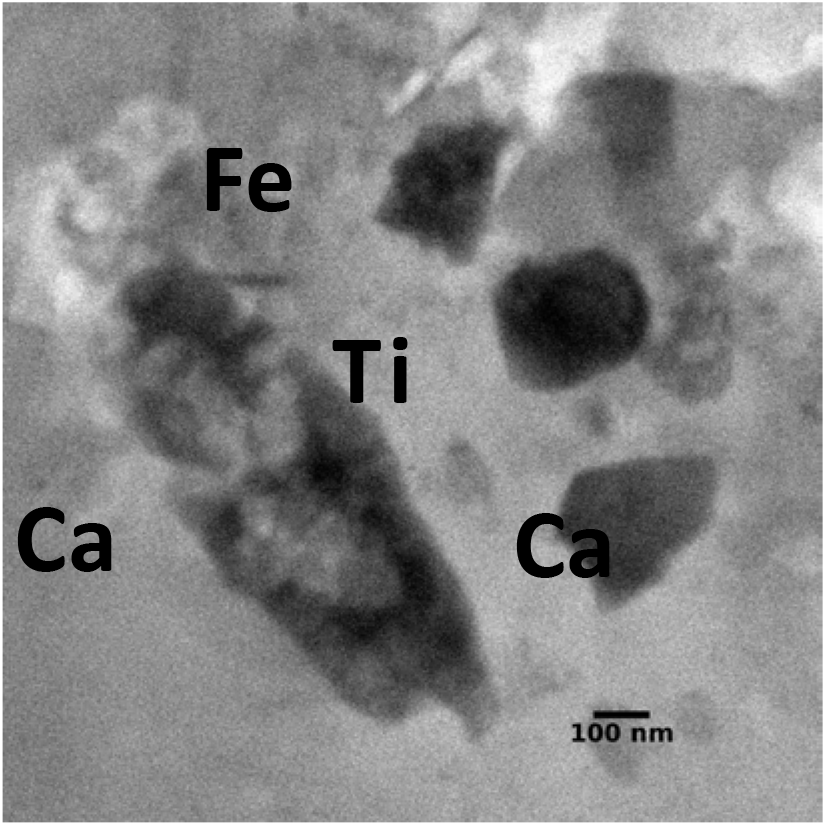
TEM showing multiple crystalline particles in the imaged field of a square of the PD stage 5 grid. The two calcium-rich crystals do not contain phosphorus; the titanium-rich crystal is protein-coated TiO_2_; the iron-rich crystal is Fe(O)OH.

## Results

Submicron crystalline particles were observed in all four PD *substantia nigra* specimens. The 3 squares-search of the PD stage 1 grid yielded one crystalline COD ghost, 3 crystalline TiO_2_ particles and 3 crystalline iron oxide particles; the 3 squares-search of the PD stage 2 grid yielded one crystalline COD ghost, 4 crystalline TiO_2_ particles and 2 crystalline iron oxide particles; the single-square search of the PD stage 4 grid yielded one crystalline TiO_2_ particle and 2 crystalline iron oxide particles; and the search of only a part of a single square of the PD stage 5 grid yielded 3 COD ghosts, 4 TiO_2_ and 2 iron oxide, for a total of 5 COD, 12 TiO_2_ and 9 Fe(O)OH particles. Of the 4 x 314 = 1,256 squares of the 4 grids 7 were fully searched and one was partially searched, meaning that the fraction of the searched squares was about 7.5/1,256=0.006. With about 0.01 of the volume of the dispersions having been applied to 4 grids, the fraction of the observed crystals was 6 x 10^−5^, meaning that each observed crystal represented about 1.6 x 10^4^ crystals in the 4 x 5 mg = 20 mg total tissue sampled, or 8 x 10^5^ crystalline particles per gram of *substantia nigra.* Because we find > 5 COD, > 12 TiO_2_ and > 9 iron oxide particles, there are more than 4 x 10^6^ COD, 10^7^ TiO_2_ and 7 x 10^6^ iron oxide particles per gram of *substantia nigra.*

Figure 1 shows a TEM image of the crystals in the PD stage 5 specimen with calcium-, titanium-, and iron-rich crystalline particles in the same field. The calcium-containing crystalline particles, an example of which is seen in Figure 2, did not contain phosphorus above the background-level, ruling out their being calcium phosphate, apatite or calcium pyrophosphate crystals. In the 10^−8^ torr vacuum of the chamber of the transmission electron microscope the imaged COD crystals promptly pulverized as they are dehydrated, then gradually lost CO_2_ and CO approaching, when completely de-gassed, the expected 71 weight % of CaO, their electron diffraction patterns changing while they are de-gassed. (Figure 2)

**Figure 2.**
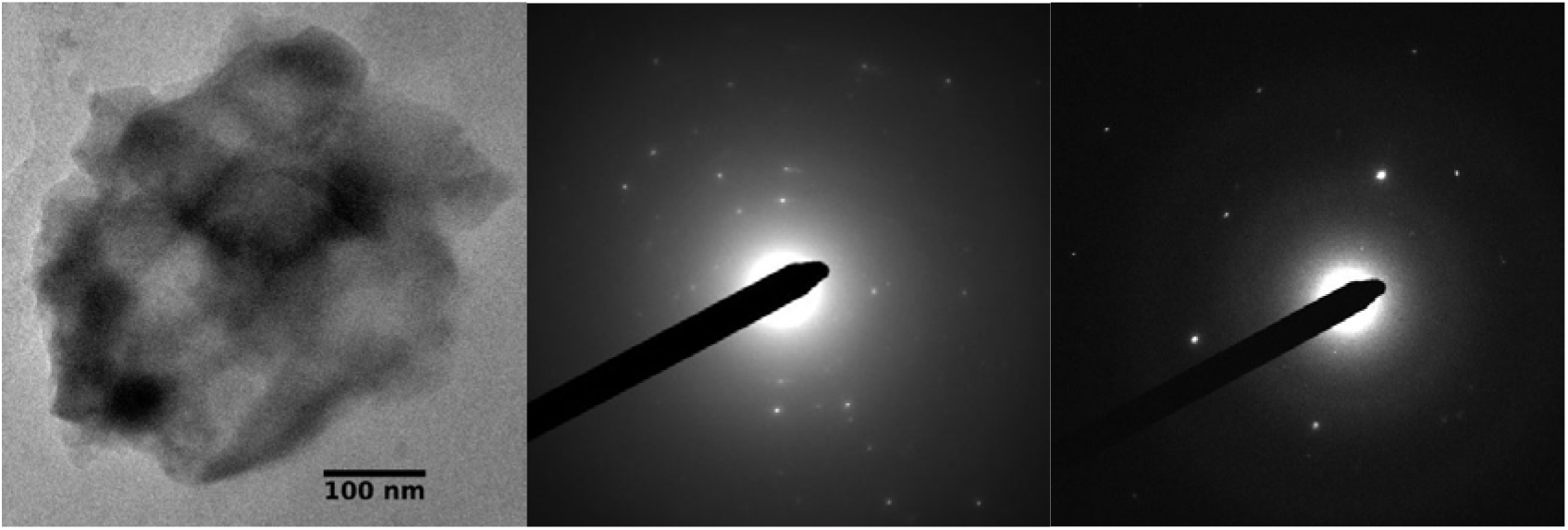
*Left.* Ghost of a COD particle of the PD stage 5 *substantia nigra* containing 53 ± 5 weight % calcium (excl. carbon), increasing as the particle decomposes under the electron-beam to 64 ± 5 weight %. *Center and right:* Changing electron diffraction patterns of the unstable particle when heated by e-beam under the TEM chamber’s 10^−8^ torr vacuum.

Figure 3 shows crystalline TiO_2_ particles of PD stages 1,2, 4 and 5 *substantia nigra.* Their 42 weight % −54 weight % titanium content is less than the expected 59.5 weight % for TiO_2_, implying that they are protein-coated. The difference between the observed 46.5 average weight % TiO_2_ and the expected 59.5 % is consistent with protein constituting about 20 % of the mass of the particles. As seen in Figure 3, the adsorbed protein is thick enough to round the all edges and corners of all the face-sharing mono-crystals constituting the particles.

**Figure 3.**
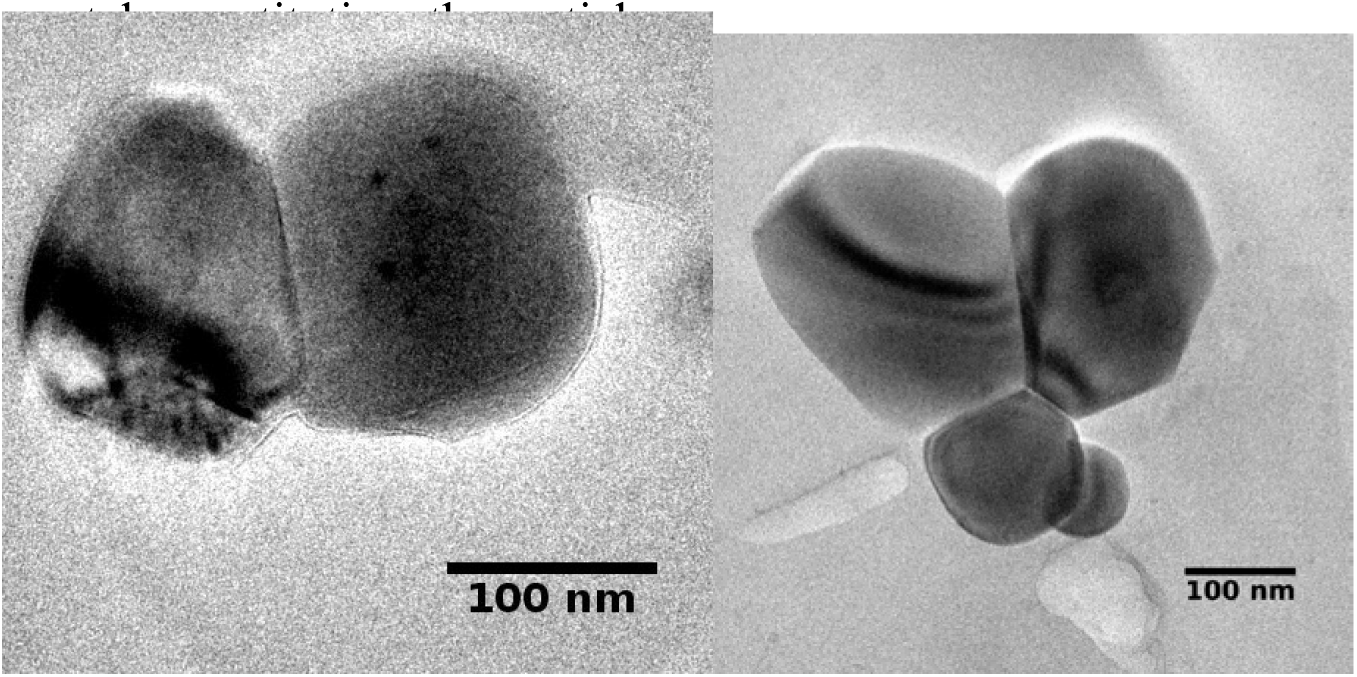
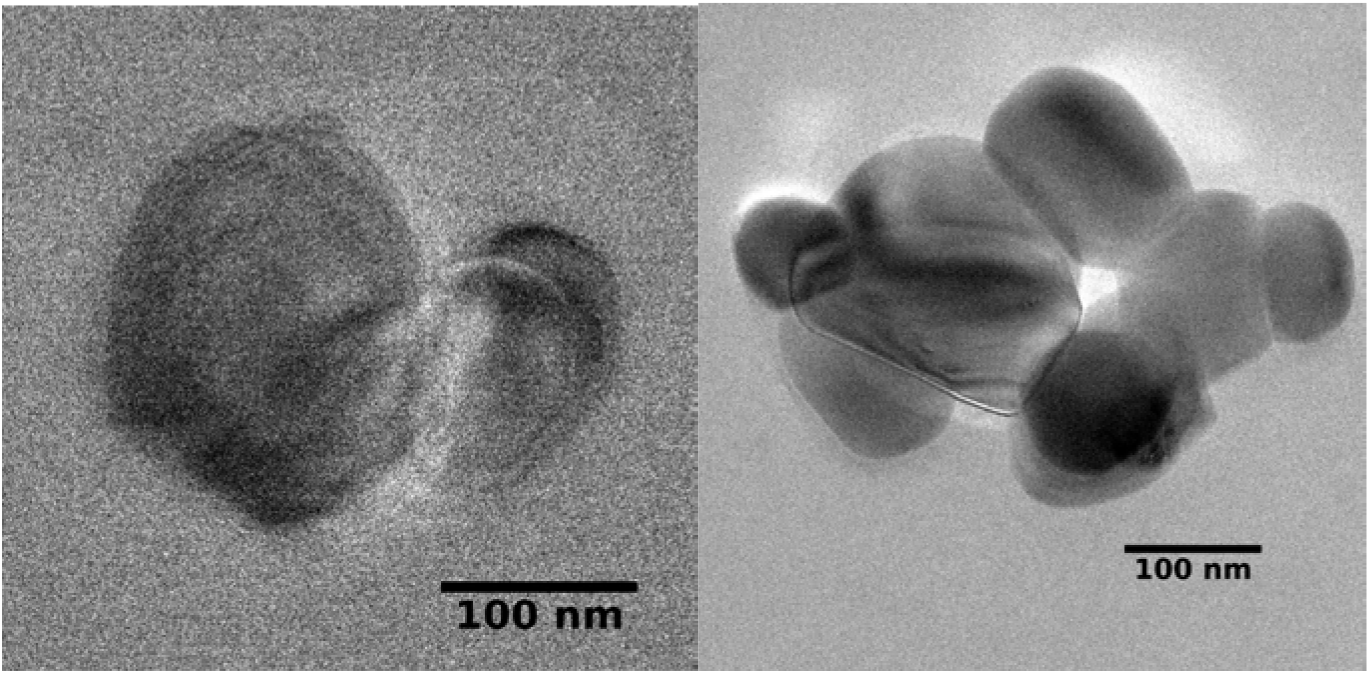
Crystalline TiO_2_ particles of the PD *substantia nigra. Top left*, from the PD stage 1 specimen, containing 54 ± 5 weight % titanium; *Top right*, from the PD stage 2 specimen, containing 42 ± 5 weight % titanium; *Bottom left*, from the PD stage 4 specimen, containing 46 ± 5 weight % titanium; Bottom right, from the PD stage 5 specimen, containing 44 ± 5 weight % titanium. The expected weight % of titanium in TiO_2_ is 59.5 %, implying that all particles are protein-coated. Protein-coating is also evident from the smoothing of the edges and corners of the crystals.

Figure 4 shows crystalline iron oxide particles of the PD stages 1,2 and 4 *substantia nigra* grids. Compositions of two are consistent with the 63 weight % iron content of Fe(O)OH, the mineral Goethite; the third shows only 51 ± 5 weight % iron content, attributed to the particle being protein-coated. Biogenic iron oxide crystals are known to abound in the human brain. [63, 64] The average iron content of 57.6 ± 5 % suggests, but does not establish, that the crystals are coated with protein constituting less than 10 % of the particle-mass.

**Figure 4.**
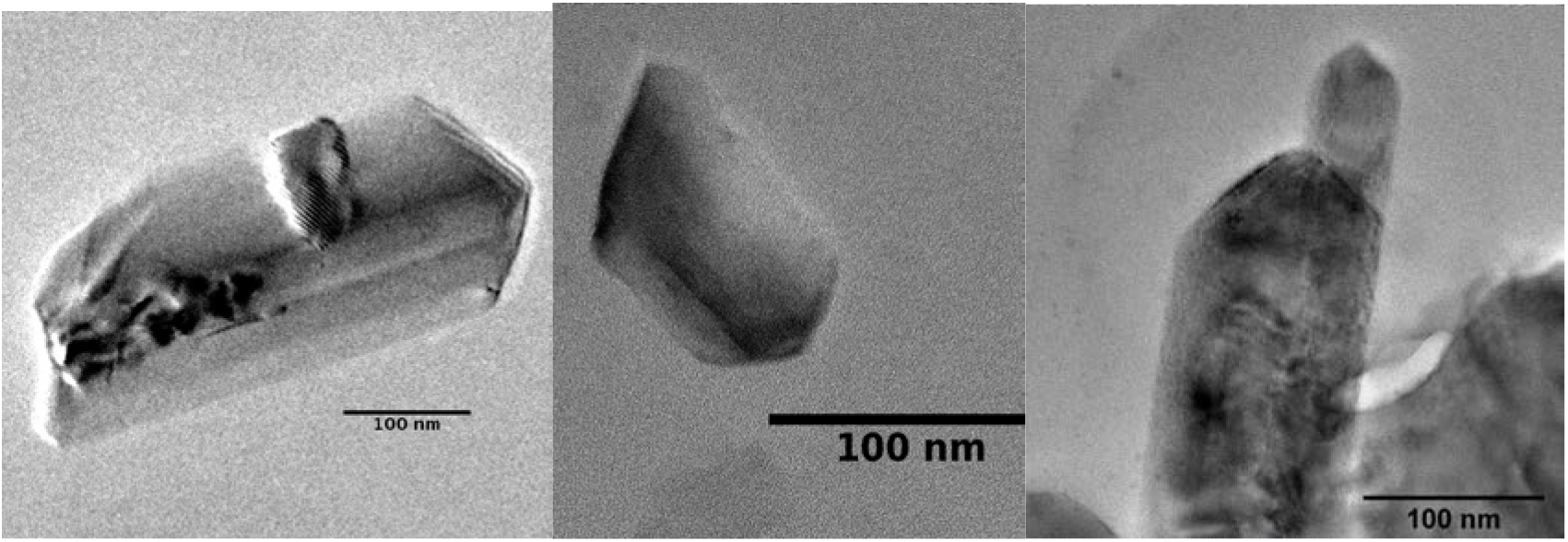
Crystalline iron oxide particles of the PD *substantia nigra. Left*, from the PD stage 1 specimen, containing 61 ± 5 weight % iron consistent with the expected 63.6 weight % in Goethite, Fe(O)OH. *Center*, from the PD stage 2 specimen, containing 51 ± 5 weight % Fe. The less than theoretical weight % of iron implies that the crystal is protein-coated. *Right*, from the PD stage 4 specimen, Fe-content 61 ±5 weight %, consistent with the expected for Goethite, Fe(O)OH. The crystals exhibit both sharp and rounded corners and edges implying that only some adsorb a thick protein layer.

## Discussion

The results show that the human PD *substantia nigra* contains submicron-sized crystalline COD and TiO_2_ particles. Both are phagocytized, [25, 35–39] the TiO_2_ also by microglia [20, 40]. In mice, activation of microglial NLRP3 inflammasome by misfolded α-synuclein causes the death of proximal dopaminergic neurons. [1, 3, 8–11] The found TiO_2_ particles are coated with a protein-layer, constituting about 20 % of their mass. TiO_2_ shown to adsorb and orient peptides and polymers [27–29] could also adsorb α-synuclein or its precursor peptides. It remains to be seen if the protein-layer found on the TiO_2_ crystals is or comprises α-synuclein and if it does, whether the α-synuclein coating is misfolded.

In humans COD inflammasome-activation causes nephropathy. [24] Just as the phagocytized α-synuclein activates the microglial NLRP3 inflammasome. Both COD and TiO_2_ are phagocytized and NLRP3 inflammasome activating. [20, 24, 25, 38, 41–47] If they activate also the microglial NLRP3 inflammasome, as is likely, proximal dopaminergic neurons would die. Because the number of microglia approaches the number of neuronal cells of the brain [65] each dopaminergic cell has on the average more than one proximal microglial cell. Consequently, in absence of segregation of the dopaminergic substantia nigra cells and the glia, death of the majority of the dopaminergic cells is expected when the NLRP3 inflammasome is activated in a large fraction of the glia. The human brain weighs about 1.5 kg and the count of its microglia is about 5 x 10^10^[65], meaning that there about 3 x 10^7^ microglia per gram. Considering that we find > 4 x 10^6^ COD and > 9.6 x10^6^ TiO_2_ particles per gram of *substantia nigra*, the number-density of particles suffices for the killing of a large fraction the dopaminergic cells.

Hydrated calcium oxalate precipitates when the solubility product of the concentrations of Ca^2+^ and oxalate is exceeded, that is when the Ca^2^+ concentration is very high, or the oxalate concentration is high, or when both concentrations are moderately elevated. COD can precipitate if Ca^2^+ stored in the endoplasmic reticulum is released into its cytoplasm, or if oxalate is excessively formed from ascorbate or citrate. Of the two, ascorbate is the more likely precursor, because in stimulated murine macrophages its concentration is about 100-fold higher than it is in the extracellular fluid, reaching 10 mM. [66] It is, therefore not surprising that a massive ascorbic acid overdose caused lethal calcium oxalate nephropathy in a patient.[67] In the ascorbate pathway, part of the ascorbate is oxidized to dehydroascorbate and is converted to 2,3-diketogluconate. 2,3-diketogluconate is oxidized by H_2_O_2_ to oxalate. [68] Oxalate production is pronounced in inflammatory M1 macrophages, but not in non-inflammatory M2 macrophages [69]; such may be the case also in stimulated microglia. Because COD stimulates macrophages. [37] An amplified inflammatory feedback loop results when more 2,3-diketogluconate-oxidizing H_2_O_2_ is produced by repeated O2-oxidation of the ascorbate-reduced pre-melanin, the pre-melamine being produced by O2-oxidation of dopamine. [70] The loop is further amplified by activation of the glia by the formed COD.

Titanium compounds have no known role in human physiology, although implanted titanium alloys do produce erosion products found in the marrow[71] and result in measurable plasma-concentrations of titanium compounds.[72] Reviews of the toxicology of titanium focus, however, on ingested and inhaled white TiO_2_ pigment particles, [61, 73–75] widely used as white pigments in foods and medications. The ingested pigment particles pass into the bloodstream[76] and are transported to the human liver, [77] spleen[77] and pancreas[78]. In animals inhaling the pigment, TiO_2_ particles were detected also in the brain.[61] Although the TiO_2_ particles are phagocytized and activate the NLRP3 inflammasome [20, 41, 47, 79, 80] their human pathogenicity is less well established than that of the COD particles.[37, 69, 81] Nevertheless, treatment of dopaminergic neurons with TiO_2_ has been shown to increase α-synuclein expression and to cause its aggregation in a dose-dependent manner. [26] Future studies might show if the observed thick protein-coating of the TiO_2_ particles (Figure 3) is the misfolded α-synuclein of Lewy-bodies. If this would be so, TiO_2_ might template the PD-underlying misfolding of this PD-hallmark protein.

Biogenic submicron iron oxide particles are well known residents of the human brain [63, 82] In the rat, brain nasally-instilled iron oxide damages dopaminergic neurons. [83] There are, however, no reports establishing that iron oxide activates the NLRP3 inflammasome. Siderosis, resulting of inhalation of iron oxide crystals by miners and steelworkers, is a relatively mild and reversible disease.

In conclusion, should future studies establish that the crystalline COD particles that were found in the *substantia nigra* cause PD, mitigation by long-known and inexpensive means of reducing COD-caused urolithiasis, such avoidance of hypomagnesemia, [84–87] could be considered. Should future studies establish that TiO_2_ particles cause PD, alternative white foods and medication colorants should be considered.

## Acknowledgement

The authors thank Dr. Karalee Jarvis for her expert guidance of the transmission electron microscopy.

